# The give and take in photosymbiosis unraveled by metabolomics of Radiolaria

**DOI:** 10.1101/2025.02.14.638264

**Authors:** Vera Nikitashina, Benjamin Bartels, Joost Samir Mansour, Charlotte LeKieffre, Johan Decelle, Christian Hertweck, Fabrice Not, Georg Pohnert

## Abstract

Marine planktonic Radiolaria harboring symbiotic microalgae are ubiquitous in the oceans and abundant in oligotrophic areas. In these low-nutrient environments they are among the most important primary producers. Systematic studies of radiolarian biology are limited because Radiolaria are non-culturable and prone to damage during sampling. To obtain insight into the mechanistic basis of radiolarian photosymbiosis we address here the metabolic contributions of the partners to the performance of the holobiont. Therefore, we describe the metabolic inventory of two highly abundant photosymbiotic Radiolaria – colony-forming Collodaria and single-celled Acantharia and compare their metabolomes to metabolomes of respective free-living algae. Most of the metabolites detected in the symbiosis are not present in the free-living algae, suggesting a significant transformation of symbionts’ metabolites by the host. The metabolites identified in both the holobiont and the free-living algae encompass molecules of primary metabolism and a number of osmolytes, including dimethylsulfoniopropionate. Mass spectrometry imaging revealed the presence of dimethylsulfoniopropionate in both the symbionts and host cells, indicating that the algae provide osmolytic protection to the host. Furthermore, our findings suggest a possible dependence of Collodaria on symbiotic vitamin B_3_. Distinctive differences in phospholipid composition between free-living and symbiotic stages indicate that the algal cell membrane may undergo rearrangement in the symbiosis. Our results demonstrate a strong interdependence and rewiring of the algal metabolism underlying Radiolaria-microalgae photosymbioses.

## Introduction

The symbiotic association of photosynthetic and heterotrophic organisms, is widespread in both terrestrial and aquatic ecosystems [1–3]. This interaction is a crucial biological process for the ecology of microbial communities, as it governs nutrient and element cycles. Additionally, photosymbiosis played a key role in evolution on Earth, with the emergence of photosynthetic eukaryotes resulting from the symbiosis between cyanobacteria and non-photosynthesizing organisms [4].

In marine ecosystems, photosymbiosis is mainly studied in corals, as understanding of this association is a key to gaining insight into coral bleaching events. These events crucially affect marine ecosystems and are closely linked to the fate of corals and their symbionts [5, 6]. Planktonic photosymbioses have received much less attention, despite their important contribution to major biogeochemical processes and their broad taxonomic diversity [7, 8]. In particular, planktonic Radiolaria host various classes of microalgal symbionts and are abundant and widely distributed in the ocean [9]. Therefore, they play important roles in the global carbon fixation and flux to the deep ocean, oxygen production, and contribute significantly to the marine biomass [8, 10]. Radiolaria are very fragile and easily damaged during bulk sampling, and they have not yet been successfully cultured. Accordingly, our knowledge about these organisms and the mechanisms underlying their photosymbiotic interactions is limited.

Given the impact of global climate change and the well-known sensitivity of coral photosymbiosis [11], we set out to investigate radiolarian photosymbioses. Our main objective was to obtain fundamental understanding of the metabolic basis of radiolarian photosymbiosis. Our study focuses on two ubiquitous groups of photosymbiotic Radiolaria: the colony forming Collodaria and the Acantharia (Fig. 1). The main symbiont of Collodaria is the dinoflagellate *Brandtodinium nutricula* [12], whereas the dominant symbionts of Acantharia belong to the haptophyte genus *Phaeocystis* [13, 14]. The photosymbiotic relationships of Radiolaria are believed to be obligate for the host, as Collodaria survive longer when exposed to light, in comparison to dark controls [15]. The symbionts of Collodaria can be isolated and grown in culture as a free-living stage, indicating a facultative symbiotic relationship for the algae [12]. In contrast, the acantharian symbiont cannot be isolated and cultured after the symbiotic stage, despite much effort [13]. Molecular evidence based on genetic markers indicates that symbionts are acquired from the environment from extensive free-living populations [13, 14]. Inside the host, the cell division of the symbiont is blocked, and its morphology changes. Particularly enlarged plastids, in comparison to those of free-living *Phaeocystis*, are observed and suggest a boosted photosynthetic activity. [13, 16].

**Figure 1:**
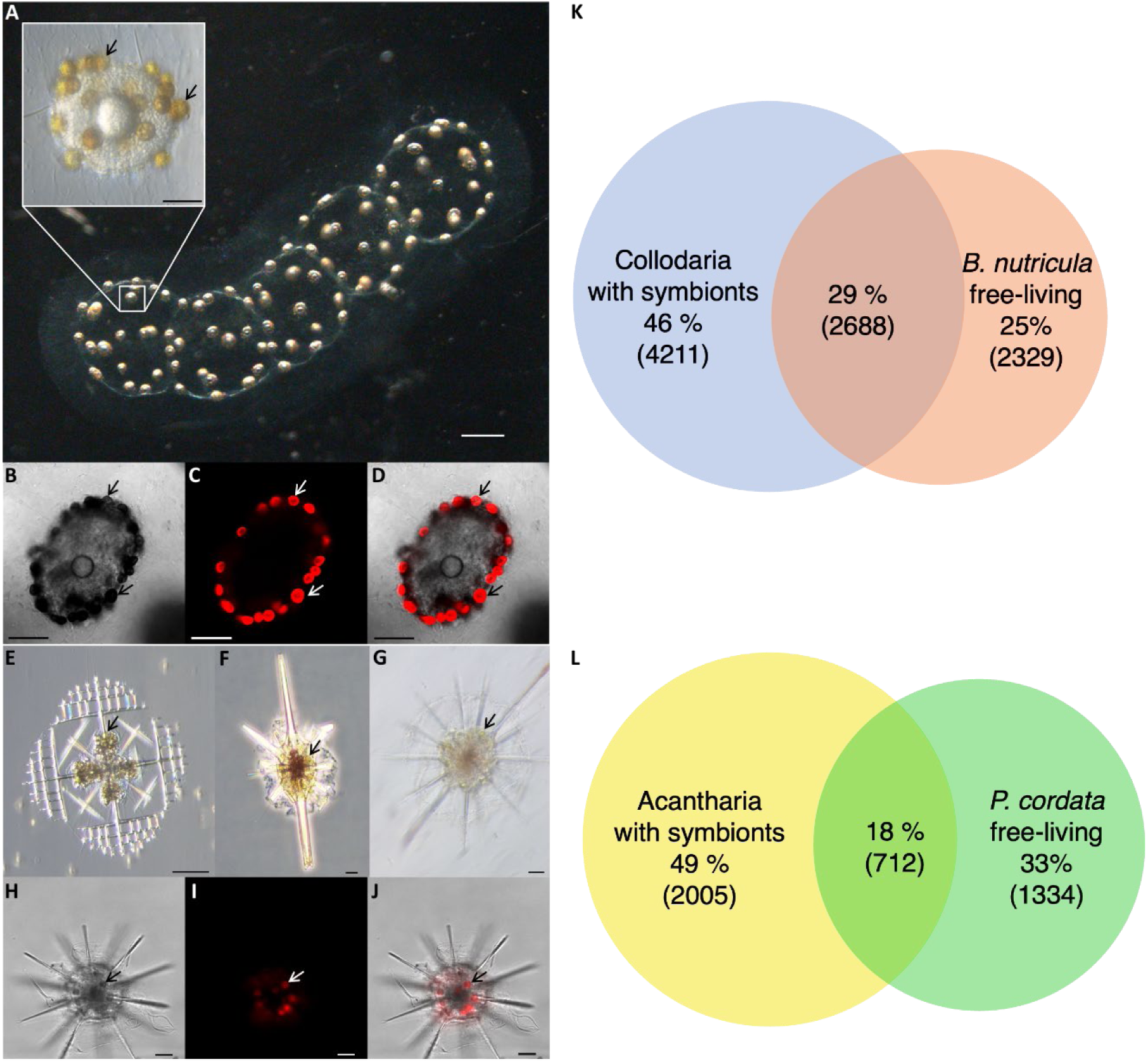
Symbiont-bearing Radiolaria and their metabolic composition in comparison to their free-living symbionts. **a** Colony of Collodaria under a brightfield microscope, the insert shows a collodarian central capsule at higher magnification (brightfield differential interference contrast microscopy); **b** – **d** Collodarian central capsule (**b**) with chlorophyll *a* autofluorescence signals (**c**) and overlay (**d**) under a confocal microscope. The arrows point to single algal cells; **e** – **g** different Acantharia cells under a brightfield differential interference contrast microscope, **h** – **j** a single Acantharia cell (**h**) with chlorophyll *a* autofluorescence (**i**) and overlay (**j**) under a light-sheet microscope; scale bars represent 500 µm for **a**, 50 µm for the insert and **b**-**d**, 20 µm for **e** – **j**. Venn diagrams for features from metabolomic comparisons detected in symbiont-bearing Radiolaria, Collodaria (blue) and free-living *B. nutricula* (red) (**k**), Acantharia (yellow) and free-living *P. cordata* (green) (**l**); the percentage of the total number of features detected is given and numbers in brackets indicate the number of features.

Metabolomics approaches have been successfully applied to describe metabolic interactions within photosymbiotic systems, for instance between corals and their photosymbionts [17]. These studies shed a light on the partners’ contributions and metabolic mechanisms of their stress responses [18–21]. Moreover, it was possible to identifying key metabolites responsible for the symbiosis stability, among others the common marine osmolyte DMSP [22, 23].

The opportunity to explore metabolic interactions within radiolarian photosymbiosis and to identify the roles of the respective partners led us to analyze metabolic composition of symbiont-bearing Radiolaria and their photosymbionts. By examining two phylogenetically and morphologically distinct photosymbiotic Radiolaria we aimed to uncover unique and shared metabolic features underlaying plankton photosymbiosis.

Here we demonstrate that the metabolic composition of symbiont-bearing Radiolaria differs from that of the free-living stage of symbionts in both studied systems. Notably, the phospholipid composition is consistently re-wired upon symbiotic association. Among the shared metabolites detected in both, symbiont-bearing Radiolaria and free-living algae, are diverse osmolytes. Using mass spectrometry imaging (MSI), we localize the distribution of metabolites in the photosymbiont-bearing Radiolaria and explain metabolic interactions underlying the radiolarian photosymbiotic system.

## Materials and methods

### Solvents

For endometabolome extraction, LC-HR-MS analysis, and analytical standard preparation (if not stated otherwise), the following solvents were used: methanol (SupraSolv, Merck, Germany), ethanol (LiChrosolv, Supelco, Merck, Germany), chloroform (HPLC grade, Fisher Scientific, UK), acetonitrile (CHEMSOLUTE, Th. Geyer, Germany), water (Chromasolv Plus for HPLC, Honeywell, Germany).

### Sampling of Radiolaria

Collodaria were collected in the bay of Villefanche-sur-Mer (France, 43°41‘10’‘N, 7°18’50‘’E) during October 2019 using a plankton net of 2 mm mesh size. Acantharia were collected at the same sampling site during September 2022 using a plankton net of 200 µm mesh size. Sampling was performed by slowly towing of the nets at the subsurface.

Collected samples were diluted into buckets with seawater freshly collected from the same sampling site. Single colonies of Collodaria and single Acantharia cells were handpicked under a stereomicroscope and transferred into filtered seawater (FSW) (0.2 µm pore size). For Collodaria, morphologically similar colonies containing symbiotic algae were isolated. For Acantharia, cells with similar morphology containing symbiotic algae were collected. Due to the demand of comparably high amounts of cells for one sample, the precise identification of individual Acantharia cells was not possible.

Isolated organisms were incubated in FSW for at least 1 h followed by transfer to FSW, this procedure was performed at least 3 times to clean organisms from attached particles. After that, the organisms were transferred into 15 mL falcon tubes (Sarstrdt AG & Co. KG, Germany) with a minimal amount of FSW, immediately frozen in liquid nitrogen, and stored below –70°C till further processing. Acantharia samples were briefly centrifuged (1,000×g, 1 min, RT) and excess water was carefully removed by pipetting before freezing. The minimal amount of FSW corresponding to the volume that was taken with the organisms was prepared as a blank. Microscopic images were taken as described in supplementary information (Appendix S3).

For Collodaria, 20 colonies per sample and 500 µL of the FSW for the blank were used, for Acantharia, 300 cells per sample and 180 µL of FSW for the blank were used.

### Strains and culture conditions of the microalgae

Strains of *Brandtodinium nutricula* RCC 3387 and *Phaeocystis cordata* RCC 1383 were obtained from the Roscoff Culture Collection (Roscoff, France). The strains were grown under sterile conditions, the associated microbiota were not removed. *B. nutricula* cultures were grown in 160 mL culture flasks containing 130 mL of the culture medium, *P. cordata* cultures were grown in 50 mL culture flasks containing 40 mL of the culture medium. Corresponding amounts of the culture medium were used as a bank. Culture medium was artificial seawater medium (ASW)[24] supplemented with an L1 culture medium kit from the Bigelow National Center for Marine Algae and Microbiota (Maine, USA) and 3 g L^−1^ sea salts (Sigma-Aldrich, USA). Stock cultures at the stationary phase were used for inoculation. Starting cell densities were 0.023×10^6^ cells mL^−1^. For the metabolome extraction, cultures of the algae were grown in triplicates till the early stationary phase.

The culturing temperature was 18 °C and the light was adjusted to 90–110 μmol photons m^−2^ s^−1^ with 14/10 light/dark cycle.

### Determination of the niacin auxotrophy for *B. nutricula* cultures

To verify whether *B. nutricula* can accumulate niacin without its presence in the medium, two sets of *B. nutricula* cultures were inoculated as described above. One set of cultures contained niacin as in the standard medium recipe, the other set of cultures was grown in medium without niacin supplementation, appropriate culture medium blanks were prepared for each set of cultures. Extraction and analysis of the samples were performed as for normal cultures of *B. nutricula*.

### Extraction

Before extraction, samples of Collodaria and Acantharia, and blanks were lyophilized. After lyophilization Collodaria samples were homogenized with a spatula, and samples containing 20 colonies were equally separated into two parts.

Algal cultures and blanks were collected on the GF/C filters (1.2 μm pore size) (GE Healthcare, US) under vacuum (750 mbar).

The extraction of the metabolome was performed as described in Vidoudez and Pohnert [25] with modifications. Briefly, 1 mL of ice-cold extraction mix (methanol:ethanol:choloroform, 1:3:1, *v*:*v*:*v*) was added to the filters with algae and blank samples in 1.5 mL Eppendorf Safe-Lock tubes (Eppendorf Quality, Eppendorf AG, Germany) or to lyophilized biomass and FSW blank samples in 15 mL falcon tubes (Sarstrdt AG & Co. KG, Germany). Samples were vortexed for 1 min, and disrupted with ultrasonication in an ultrasonic cleaner USC1200TH (VWR, Malasia) for 10 min. After this, samples were frozen in liquid nitrogen and thawed on ice. Subsequently, samples were vortexed for 1 min and sonicated for 10 min (repeated 2 times). All samples were transferred into new Eppendorf Safe-Lock tubes, algal samples and culture medium blanks were transferred without filters. Cell debris was removed by centrifugation at 30,000×g at 4 °C for 15 min, the supernatant was collected into 1.5 mL glass vials. For the algal samples, volumes for LC-HR-MS sample generation were normalized according to the cell counts in the cultures. For *B. nutricula* cultures aliquots with 10^7^ cells per sample were prepared, for *P. cordata* cultures aliquots with 3 ⋅ 10^6^ cells per sample were prepared, for the media blanks the average volume of the culture’s volume was prepared. The samples were evaporated under vacuum. Dried samples were kept under argon and stored at –20 °C.

### LC-HR-MS Sample preparation

Dried samples of Collodaria and *B. nutricula* were dissolved in 65 μL of 80% methanol (*v*:*v*), vortexed for 1 min, ultrasonicated for 10 min to ensure complete dissolution, transferred into 1.5 mL Eppendorf Safe-Lock tubes and centrifuged at 30,000×g at 4 °C for 15 min to remove any particles. Supernatants were transferred into glass vials with inserts for the measurement on C18 a column. Subsequently, 10 μL of each sample were diluted in 90 μL of methanol:acetonitrile:water (5:9:1, *v*:*v*:*v*) mixture to measure on a SeQuant ZIC-HILIC column.

For the Acantharia and *P. cordata* samples, the procedure was the same, except that the analysis with both C18 and ZIC-HILIC columns was performed on one set of samples. For this, the samples were either dissolved in 65 μL of 80% methanol (*v*:*v*), or in a mixture of methanol: acetonitrile: water (5:9:1, *v*:*v*:*v*). Between two analyses, samples were dried under vacuum.

### Standards

Solutions of standards were prepared by dissolving the respective compound in an appropriate solvent (water, methanol or a mixture of methanol:acetonitrile:water (5:9:1, *v*:*v*:*v*)) with the final concentration of 1 mg mL^−1^. The list of measured standards is presented in the supplementary information (Appendix S4).

### LC-HR-MS measurements

Samples for LC-HR-MS analysis were measured with a Dionex Ultimate 3000 system coupled to a Q-Exactive Plus Orbitrap mass spectrometer (Thermo Scientific, Bremen, Germany). The measurements were performed in positive and negative modes, heated electrospray ionization was used to generate molecular ions. Two types of separation columns were used for the analysis of the samples.

For the separation of the samples on a SeQuant ZIC-HILIC column (5 μm, 200 Å, 150 × 2.1 mm, Merck, Germany), equipped with SeQuant ZIC-HILIC guard column (20 × 2.1 mm, Merck, Germany) the duration of the method was 14.5 min with an MS runtime from 0.5 min to 9 min. Eluent A consisted of water with 2% acetonitrile and 0.1% formic acid, eluent B of 90% acetonitrile with 10% water and 1 mmol L^−1^ammonium acetate. The flow rate was set to 0.6 mL min^−1^, the gradient started with 85% solvent B, which was held for 4.0 min, gradient to 0% of solvent B (4.0–5.0 min), hold time 0% B (5.0–9.0 min), gradient to 100% of B (9.0–10.0 min), hold time 100% B (10.0–12.0 min), gradient to 85% of B (12.0–12.5 min), hold time 85% B (12.5–14.5 min).

For the separation on a THERMO Accucore C18 RP column (2.6 μm, 100 × 2.1 mm, Thermo Scientific, Germany) the duration of the method was 12.0 min with MS runtime from 0.0 to 11.5. Eluent A consisted of water with 2% acetonitrile and 0.1% formic acid, eluent B of 100% acetonitrile. The flow rate was set to 0.4 mL min^−1^, the gradient started with 100% of solvent A and held for 0.2 min, followed by a linear gradient to 0% of solvent A, 100% of solvent B in 7.6 min, held at 100% solvent B for 3.0 min, in 0.1 min to 100% of solvent A which was held for equilibration at 100% of solvent A for 1.1 min.

The instrument settings can be found in the supplementary information (Appendix S6).

### LC-HR-MS data pre-processing

Raw data were preprocessed using Compound Discoverer version 3.3 (Thermo Fisher Scientific). The standard workflow (Untargeted Metabolomics with Statistics Detect Unknowns with ID Using Online Databases and mzLogic) with default settings, except that setting for Peak Rating Filter were changed to: Peak Rating Threshold – 5, number of files – 2, was applied. Peak picking and deconvolution for the Collodaria and *B. nutricula* experiment was performed separately from the Acantharia and *P. cordata* experiments. Due to high background in the first 0.6 min of measurement for data obtained from SeQuant ZIC-HILIC column separation, the analysis of the signals was performed starting from 0.6 min of the LC run.

Further analysis and identification were performed for signals of which the average peak area of the samples was at least 5 times higher than in blank samples.

### Venn diagrams

The Venn diagrams were made using the web application BioVenn [26].

### Identification of metabolites

Identification was undertaken for the 200 most intense signals of each group (algal unique compounds, holobiont unique compounds, common compounds) detected with the positive ionization mode. Results were manually checked for processing artifacts. Identification of unknowns was performed using Sirius version 5.6.3 [27] with parameters in the supplementary information (Appendix S7).

### Collodaria sectioning

Cryo-sections were obtained from Collodaria colonies embedded in M1 medium using a Leica CM1850 cryostat (Leica Mikrosysteme, Wetzlar, Germany). The sections were cut to a thickness of 60 µm using a S35 low profile blade (Leica) and captured with the anti-roll plate. It was necessary to cut at a very flat angle and space the anti-roll plate from the blade with 3 layers of aluminum foil to achieve intact sections of this very aqueous sample. The sections were thaw-mounted on ITO glass slides (Bruker) and lyophilized overnight. The dry, mounted sections were kept at –70°C and transported to the place of analysis in a desiccator shuttle, to avoid surface condensation and subsequent delocalization of metabolites.

### Mass spectrometry imaging

The MALDI-matrices CHCA, DHAP, and DHB (all Bruker, Bremen, Germany) were tested against a panel of 32 compounds, previously identified in the metabolomic experiments as described in in the supplementary information (Appendix S8).

Optical images of the dry, thawed sections were acquired with a Keyence VHX-5000 digital microscope (Keyence Deutschland GmbH, Neu_Isenburg, Germany) at 100x magnification, using a polarizer for glare removal and the integrated stitching function. Afterwards, CHCA was applied as MALDI matrix with the M3+ sprayer (HTXImaging), using the respective sprayer method described in the supplementary information (Appendix S9). MALDI-2-MSI was performed in positive ion mode, mass range *m/z* 90–1025, in the timsTOF fleX MALDI-2 mass spectrometer (Bruker) with a step size of 20 µm, 75% laser energy, and 100 shots per position at 1 kHz repetition rate. The height and position of the sample stage was adjusted manually for each sample in combination with focus adjustment, to achieve the best MALDI-2 performance.

Data analysis was performed, and spatial distributions calculated in LipoStarMSI v2.0.1 (Mass Analytica, Sant Cugat del Vallés, Spain) [28]. The target list of the previously identified compounds of interests was searched against the compound’s exact mass +/– 0.005 Da and 2 ppm error. All spatial distributions were subjected to hot spot removal (99.75 quantile) and RMS normalization.

## Results

We analyzed the metabolic compositions of Radiolaria carrying symbionts and the respective free-living microalgae to shed light on their photosymbiotic interactions. It is not possible to obtain a native metabolome of the aposymbiotic host without the metabolites of the symbionts since Radiolaria cannot survive without their symbionts [15].

### The Collodaria - *B. nutricula* symbiotic system

We analyzed the endometabolome of symbiont-bearing colony-forming Collodaria, identified as *Collozoum pelagicum*, collected from their natural environment [29]and compared it to cultures of the free-living form of the symbiotic dinoflagellate *B. nutricula*.

The comparisons of the endo-metabolomes revealed three groups of metabolites. The largest group was only present in the Collodaria holobiont and represented 46% of all detected features (Fig. 1k). Among them we identified hypoxanthine, pipecolate, 3-methyladenine, phytosphingosine and several betaines including hydroxyproline betaine, γ-aminobutyric acid betaine (GABA betaine), and sulfobetaine (Table 1). One-third of all detected features was present in both, Collodaria holobiont and free-living algae. Analysis of these metabolites revealed amino acids, betaines such as dimethylsulfoniopropionate (DMSP), gonyol, glycine betaine, proline betaine (stachydrine), homarine (N-methyl picolinic acid betaine), alanine-and β-alanine betaines (Table 1). High diversity of betaines in the holobiont and their presence in both holobiont and free-living stage of symbionts indicates their importance in maintaining the homeostasis of both, Collodaria and their symbionts.

**Table 1:**
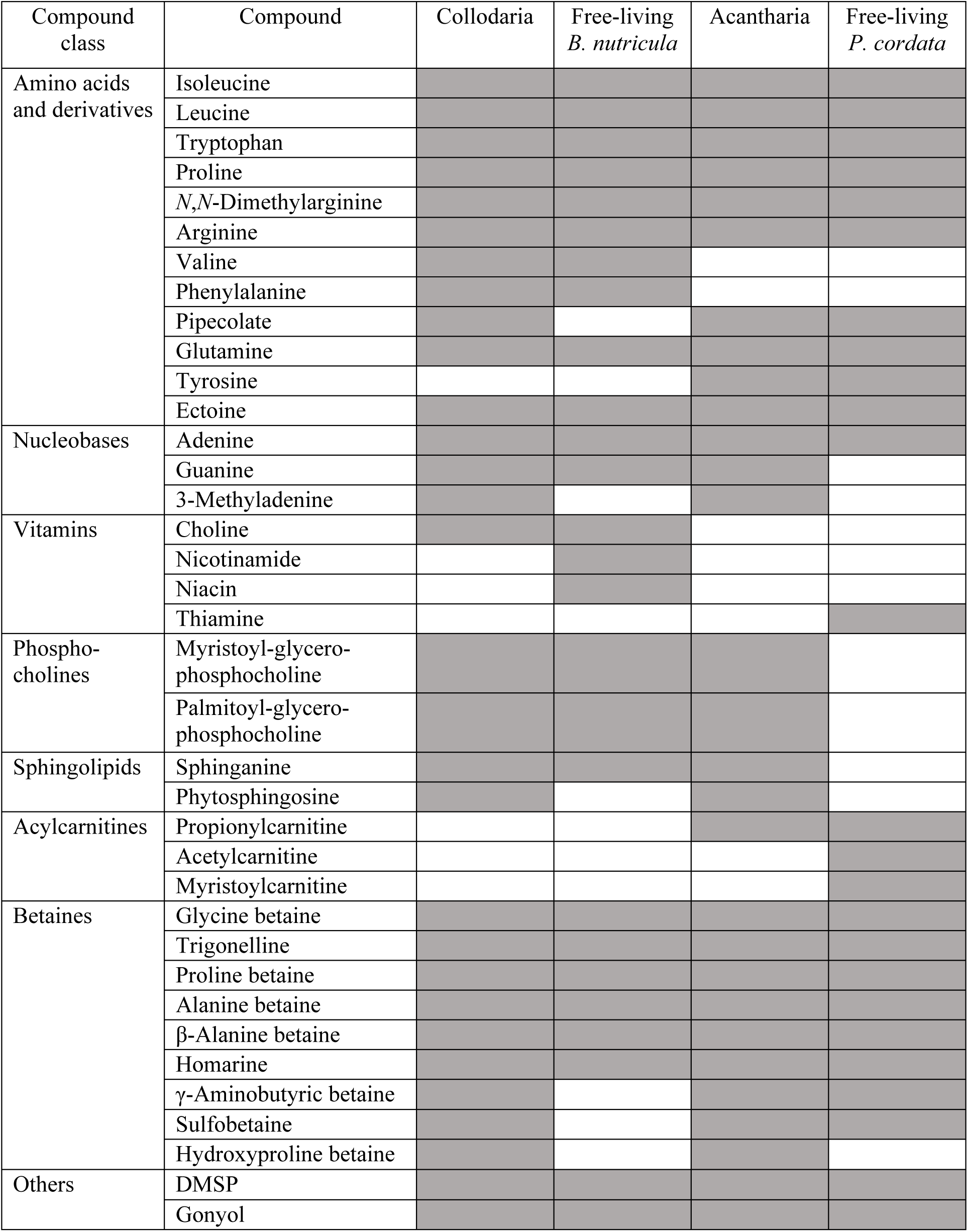
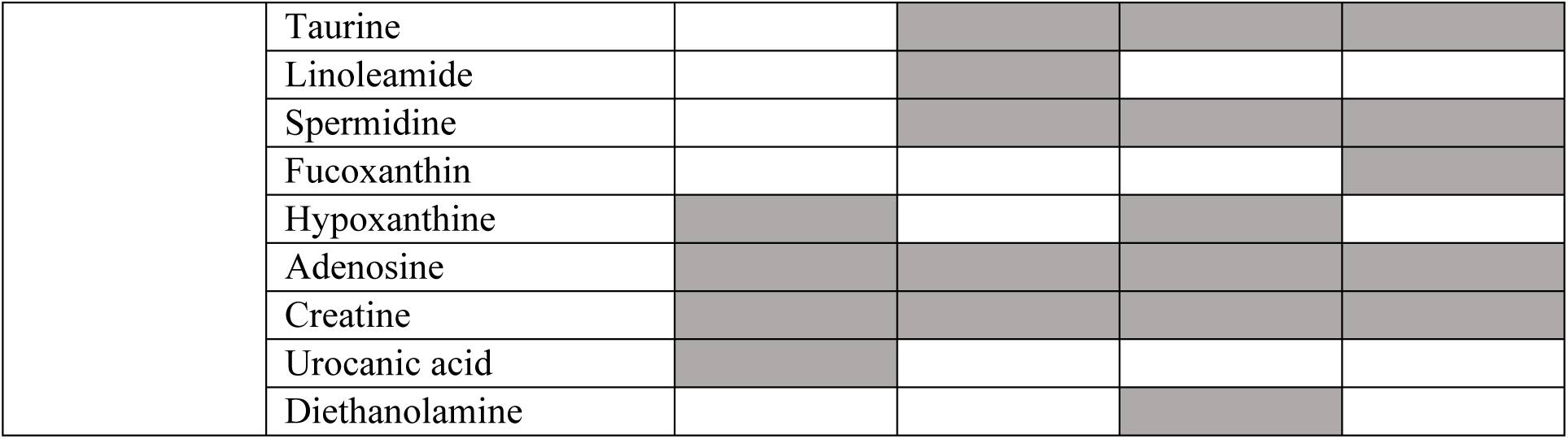
List of metabolites identified and confirmed with analytical standards in the samples of the Collodaria holobiont, free-living *B. nutricula*, the Acantharia holobiont, and free-living *P. cordata*; gray color indicates the presence of the metabolite, no color indicates absence of the compound.

The smallest group of metabolites was detected only in free-living stage of *B. nutricula* (Fig. 1k). Among these metabolites, linoleamide, taurine, and spermidine, together with two forms of vitamin B_3_ – niacin and nicotinamide, were identified. Since vitamin B_3_ might be a metabolite provided to the host by the symbionts, we tested the ability of the symbionts to grow and produce B-vitamins without niacin supplementation in the culture medium. The metabolome of algae grown with and without niacin showed that they both forms of the vitamin (niacin and nicotinamide) regardless of niacin presence in the culture medium confirming their ability to produce vitamin B_3_ (Appendix S1).

Sixteen phospholipids and their derivatives uniquely present in the endometabolome of free-living *B. nutricula* were identified to the level of compound class implying a difference in the cell membrane composition between free-living and symbiotic stages (Table 2).

**Table 2:** Putatively identified compound classes in the samples of Collodaria and free-living *B. nutricula*. The number of compounds detected from the respective class is given in brackets.

| Organism | Compound class |
| --- | --- |
| Collodaria and free-living *B. nutricula* | Ether-linked phosphatidylethanolamine (1) |
|  | Phosphatidylcholines (2) |
|  | Ether-linked phosphatidylcholines (2) |
|  | Lysophosphatidylcholines (3) |
| Collodaria | Phosphatidylcholine (1) |
|  | Lysophosphatidylcholine (1) |
| Free-living *B. nutricula* | Glycerolipid (1) |
|  | Tripeptides (3) |
|  | Dipeptides (4) |
|  | Phosphatidylethanolamines (3) |
|  | Lysophosphatidylethanolamines (4) |
|  | Lysophosphatidylcholine (4) |
|  | Phosphatidylcholines (5) |

### The Acantharia - *P. cordata* symbiotic system

In an identical approach we studied the endometabolomes of symbiont-bearing Acantharia and of the free-living *P. cordata*. Acantharia, containing endosymbionts were collected at the field site. *P. cordata* is the symbiotic microalgae of Acantharia in the Mediterranean. It has not been isolated from the host but is available as culturable strain, which was used in this study.

As for Collodaria and *B. nutricula*, we also categorized three groups of metabolites in the endometabolomes of Acantharia holobionts and cultured *P. cordata*: metabolites present in both samples, those unique to the Acantharia holobiont, and those only found in free-living *P. cordata* (Fig. 1l). Approximately half of all detected compounds were present only in the Acantharia holobiont, among them we identified hypoxanthine, 3-methyladenine, guanine, 2 sphingolipids, 2 lysophosphatidylcholines, diethanolamine, and hydroxyproline betaine (Table 1). Additionally, 28 dipeptides, identified to the compound class level, were exclusively detected in Acantharia holobionts (Table 3). The portion of metabolites detected in both samples made up less than one fifth of all detected features. Amino acids and zwitterionic compounds, such as DMSP, gonyol, and betaines were prominent among the metabolites identified from this category (Table 1). Thiamine and fucoxanthin, as well as several phospholipids identified to the compound class level, were found exclusively in free-living *P. cordata* (Tables 1 and 3). The distribution of metabolites that are common or unique for the holobiont and the free-living stage of the symbionts in the Acantharia – *P. cordata* photosymbiotic system is very similar to the Collodaria – *B. nutricula* system. This implies common mechanisms underlying Radiolaria photosymbiosis.

**Table 3:**
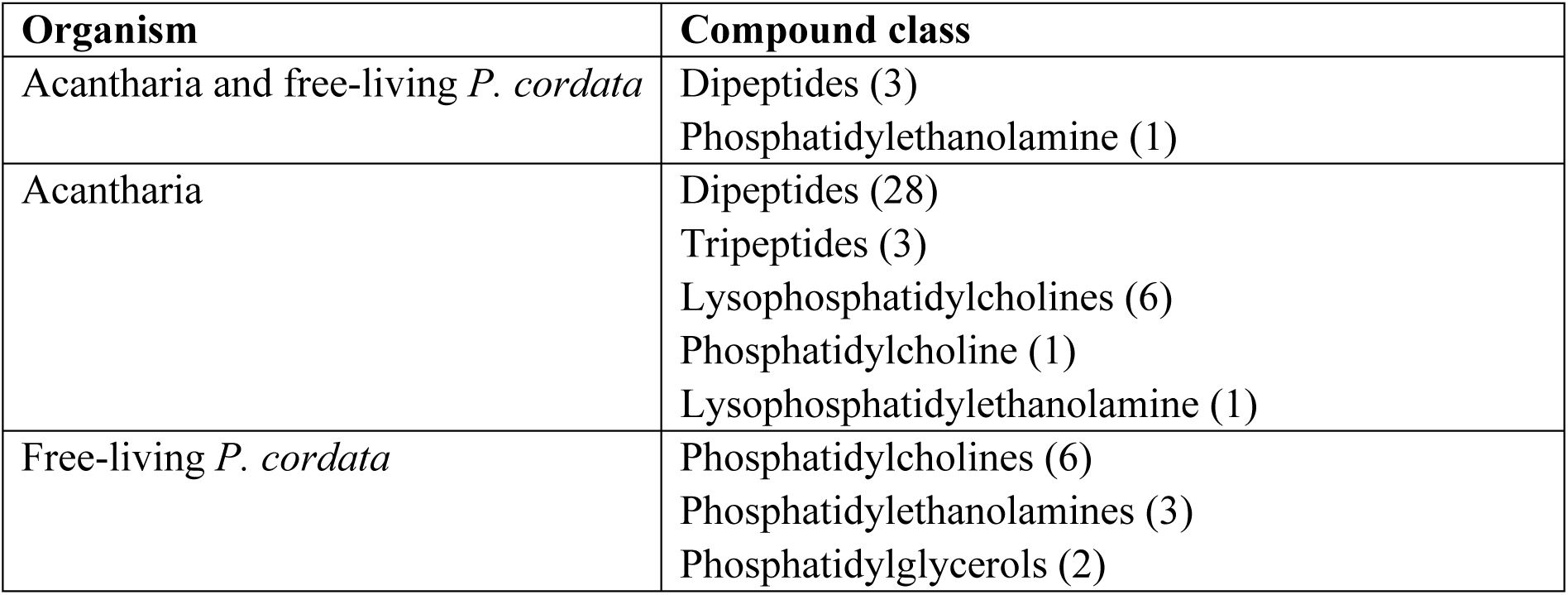
Putatively identified compound classes in the samples of Acantharia and free-living *P. cordata*. The number of the compounds from the class is given in the brackets.

### Comparison of the Collodaria - *B. nutricula* and Acantharia - *P. cordata* symbiotic system

Similar for both the Collodaria - *B. nutricula* system and Acantharia - *P. cordata* system, around half of all detected compounds were present only in holobionts. The portion of metabolites detected in both holobionts and free-living algae was higher for the Collodaria - *B. nutricula* system (one third of all detected features). In turn, the proportion of unique features for the free-living algae was higher for *P. cordata* than for *B. nutricula* – one third of all detected metabolites fell in this category for the free-living haptophyte. For *B. nutricula* and the Collodaria system, only one quarter of metabolites was unique for the algal symbiont (Fig. 1).

Most of the identified compounds were detected in all studied organisms, however there were some differences in the composition of metabolites between the two investigated symbiotic systems. Thus, five detected betaines were present in all organisms, but another two betaines – γ-aminobutyric acid betaine and sulfobetaine, were not found in the metabolome of the free-living *B. nutricula*. Whereas hydroxyproline betaine was only detected in the metabolomes of both holobionts. Another dissimilarity was observed in the presence of two lysophosphatidylcholines and sphinganine. These compounds were present for both, Collodaria holobiont and cultivated *B. nutricula*, but found only in the endometabolome of Acantharia holobiont but not in the free-living *P. cordata*. In turn, a non-proteinogenic amino acid pipecolinic acid was detected in both Acantharia and free-living *P. cordata*, and in Collodaria holobiont but not in free-living *B. nutricula* (Table 1).

A major variation between both holobionts and their free-living microalgae was in the composition of phospholipids. For each of the free-living algae more than 10 unique phospholipids were putatively assigned (Tables 2 and 3).

### Spatial distribution of metabolites in the Collodaria holobiont

We performed MSI on sections of frozen Collodaria colonies, to investigate the spatial distribution of metabolites in the holobionts. During data analysis, we focused on the masses of compounds previously identified in the metabolome of Collodaria holobionts and free-living *B. nutricula* (Table 1).

Several *m*/*z*-values with distinctive spatial distributions in areas of symbionts, as well as Collodaria central capsules and areas connecting them, were putatively assigned to ectoine, proline, glutamine, gonyol, and creatine as the observed *m*/*z*-values fit the expected masses (+/– 0.005 Da and 2 ppm error). In the outer regions of the colony signals for ectoine, proline, tryptophane, glutamine, gonyol, and stachydrine were detected (Fig. 2, Appendix S2). The spatial distribution of *m*/*z*-= 135.0474, putatively assigned to the [M + H]^+^ of DMSP, exhibited high intensities in the areas of the central capsules and their connections, and outer region of the colony. Of note, the detected signal intensity of this ion was low (S/N<4). The putative assignment was performed based on its spatial distribution and the observed accurate mass. The *m/z*-values fitting homarine and trigonelline, isobaric compounds indistinguishable by MS, also exhibited a spatial distribution that correlated with areas belonging to both host and symbiont cells (Appendix S2). A different pattern of spatial distribution was observed for the nucleobases adenine and guanine. Ions with fitting *m*/*z*-values of their respective adduct ions were found in regions of Collodaria central capsules (Fig. 2, Appendix S2). A similar pattern was observed for an *m*/*z*-value corresponding to the palmitoyl-glycero-phosphocholine (Appendix S2). Urocanic acid and γ-aminobutyric acid betaine are compounds that were detected only in Collodaria colony samples, but not in free-living *B. nutricula*. In accordance, the spatial distribution of the respective ion’s *m*/*z*-values corresponded to the Collodaria central capsules and areas connecting them (Appendix S2). The presence of zwitterionic compounds in host cells and their distribution through the colony supports the suggestion that these compounds play an important role in the homeostasis of the holobiont.

**Figure 2:**
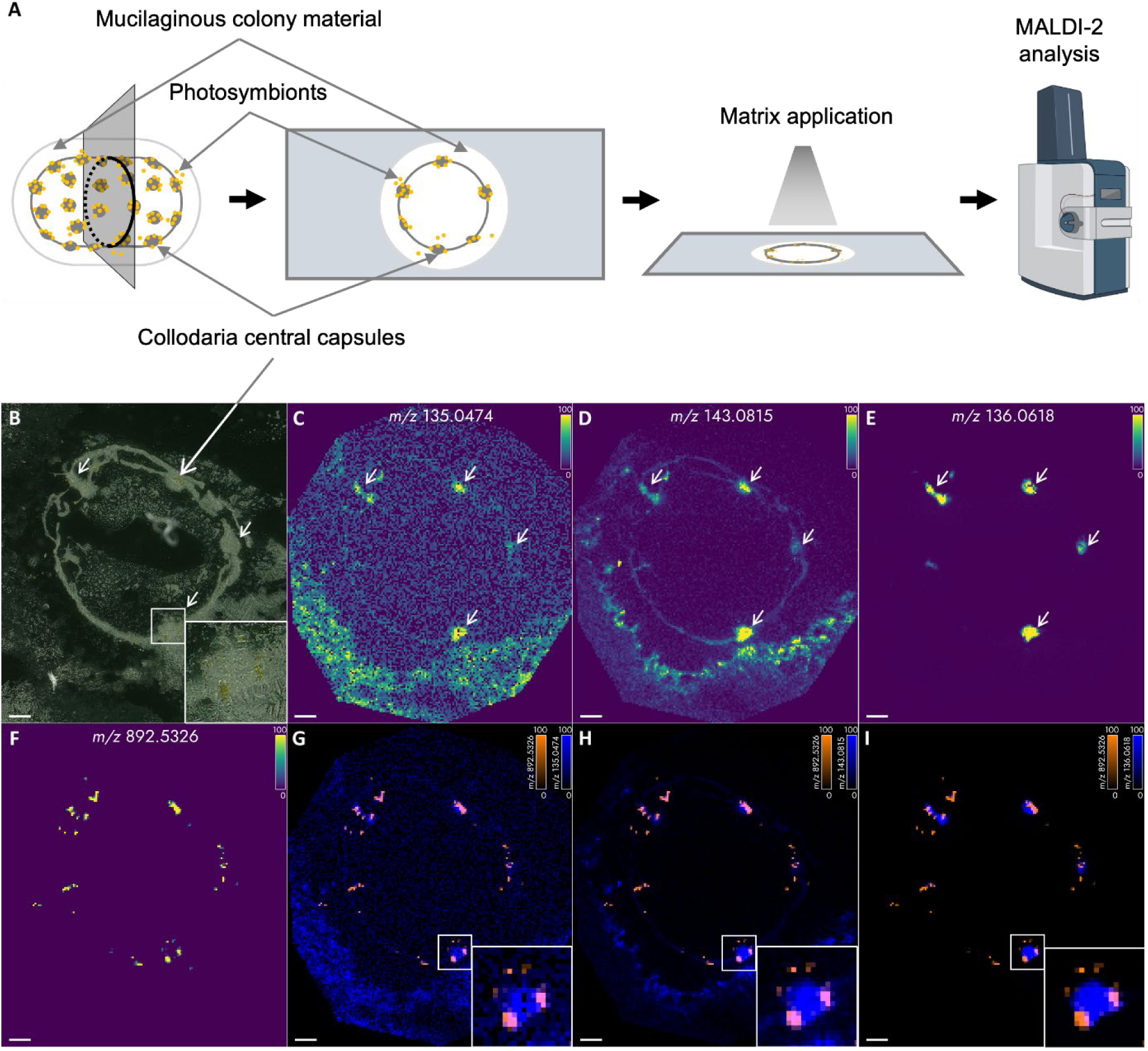
MALDI-2-MSI of a Collodaria colony section. **a** schematic workflow of Collodaria sample preparation for MALDI-2-MSI; **b** optical image of the section before analysis, white arrows indicate positions of Collodaria central capsules; **c, d, e, f** spatial distributions of *m/z* 135.0474, 143.0815, 136.0618, and 892.5326, putatively assigned to the [M+H]^+^ of DMSP, ectoine, adenine, and the [M]^+•^ of chlorophyll *a*, respectively; **g**–**i** overlay of **c**–**e** with the Chl *a* signal, indicating the position of algae cells. The scale bar represents 200 µm in all panels. The timsTOF instrument image is from BioRender.com

## Discussion

Radiolaria are dependent on microalgae but since they are unculturable there is a substantial lack of knowledge about the metabolic mechanisms underlying the photosymbiotic relationships. In a system comparison based on holobionts and free-living symbionts we derive a model for the metabolic interdependence. This will contribute to the prediction of responses of marine ecosystem to the global climate change and anthropogenic interventions.

Despite mechanistic and taxonomic differences in the symbiotic relationships between the Collodaria and *B. nutricula* and the Acantharia and *P. cordata*, metabolites unique for the holobiont represented the largest fraction of identified compounds. Such high proportions of unique metabolites in the holobionts imply a complex interdependent metabolism within the symbiosis. Also, it indicates a pronounced transformation of algal metabolites by the host and/or the induction of metabolite biosynthesis by the symbionts during association. An example of a holobiont unique metabolite is hypoxanthine, which was detected in both Collodaria and Acantharia holobionts but not in the free-living algae. Hypoxanthine can play a role as a nitrogen reservoir [30] that is accumulated by the holobionts as a storage form of nitrogen while living in a nutrient replete environment and that can be mobilized under nutrient limitation.

Metabolites common for holobiont and free-living algae, in the symbiotic systems can exclusively originate from the symbionts. Alternatively, both, host and algae can contribute to their production, as it was shown for coral photosymbiosis [22]. Most metabolites identified within this group for both radiolarian symbiotic systems were molecules of primary metabolism, and particularly amino acids such as arginine. Arginine can contribute up to 50% of the total nitrogen storage in seeds and embryos of several plants but also accumulates substantially in algae under nitrogen replete conditions [31, 32]. Endometabolomes of both, the free-living algae and the holobiont contained comparatively high amounts of arginine, whereas MSI on Collodaria colonies showed a presence of arginine mainly in the areas of the central capsules of the host. This is in accordance with a mechanism, where in symbiosis the metabolite is transferred from the algae to the host (Appendix S2).

Moreover, we identified several osmolytes including the amino acids proline and ectoine, sulfur containing molecules such as DMSP and gonyol, and other betaines as common metabolites in holobionts and free-living algae. The osmoprotective role of these compounds was previously reported in different organisms including microalgae [33–37]. Moreover, photoprotective and reactive oxygen species (ROS)-protective roles of several betaines were proposed for the photosynthetic systems of corals’ symbionts [34, 38]. Comparatively high amounts and diversity of osmolytes in photosymbiotic Radiolaria and free-living microalgae could be required to maintain the homeostasis of the holobionts in the changing environment of the open ocean. The presence of these metabolites in the central capsules and the entire Collodaria colony revealed by MSI supports this hypothesis.

The non-proteinogenic amino acid ectoine and is widely distributed in marine microorganisms [39–43]. Since it was detected in the free-living stage of both, *B. nutricula* and *P. cordata*, it is likely that the symbionts are synthetizing ectoine in the photosymbiotic association. However, within the holobiont, this compound is no longer associated only with the algae. The spatial distribution of the respective *m*/*z*-values correlates with the Collodaria central capsules and colony matrix and is not always overlapping with the regions associated with the algae (Fig. 2). These results indicate that algae, which are internalized and likely do not experience osmotic stress anymore, provide the compound to the Radiolaria. However, the ability of Radiolaria to produce ectoine or obtain it from associated bacteria also cannot be excluded and requires further analysis. Similar spatial distribution was observed for the sulfur-containing compound DMSP. This molecule is an abundant metabolite of marine microalgae which plays a crucial role in the global sulfur cycle [44]. In some phytoplankton species, it has been estimated that DMSP can contain as much as 50% of the total sulfur content of the organisms [45, 46]. Moreover, DMSP plays an important role in the symbiosis of corals and their stress responses [23], where it can be produced not only by the symbiotic microalgae but also by non-symbiotic juvenile corals implying the contribution of the host to the common pool of DMSP [22]. Our MSI analysis located the *m*/*z*-signals for DMSP in both, symbionts and host cells, which can imply a contribution of the host to the DMSP pool of the photosymbiotic system (Fig. 2). This hypothesis is supported by the previous observations that the quantitative amount of DMSP is much higher in host - microalgae associations compared to microalgae alone [13, 47].

Compounds that were detected only in free-living algae can be explained by a reversible or irreversible change of the metabolism upon association to the host [13, 16, 48]. Alternatively, these metabolites could be further produced by the algae and metabolized by the host. The proportion of metabolites uniquely detected in free-living algae was higher in *P. cordata* in comparison to *B. nutricula*. This might result from more pronounced physiological changes of the *P. cordata* in symbiosis compared to the free-living stages [48]. This is supported by the observation of drastic morphological transformation of the symbionts upon association with the host and the impossibility to isolate it in culture [13]. In contrast, *B. nutricula*, which showed fewer unique metabolites, can be isolated from the host and maintained in culture [12]. The main metabolic difference between the holobiont and the free-living microalgae was in the composition of phospholipids for both symbiotic systems. As in most living beings, phospholipids are essential components of membranes in dinoflagellates and haptophytes[49–51]. The difference in phospholipid compositions between the holobiont and free-living algae implies not only distinct cell membrane composition of the host and the symbiont, but also a difference in the cell membrane composition between the free-living and the symbiotic stage of the algae. This is in accordance with the fact that for both, the dinoflagellate *B. nutricula* and the haptophyte *P. cordata*, morphological differences between free-living and symbiotic stages were described [12, 16, 48].

Other compounds which were detected only in free-living *B. nutricula* were two forms of vitamin B_3_ – niacin and nicotinamide. Since vitamin B_3_ is an essential nutrient [52], it should be present in all organisms, and its absence for Collodaria, Acantharia and *P. cordata* could be caused by its rapid turnover by the organisms, resulting in undetectable amounts. Higher amounts of the vitamin in *B. nutricula* cultures might be explained by the fact that the algae accumulate the vitamin without consumption by partner organisms. When in symbiosis, it provides it to the host. The ability of dinoflagellates to supply symbiotic partners with niacin was previously described in symbioses with bacteria [53, 54].

Limited knowledge about Radiolarian physiology and biochemistry, and absence of sequenced genomes hinder the elucidation of the observed interdependence with algae. Our use of metabolomics combined with MSI allows to, nevertheless, gain an insight into metabolic re-wiring of the algae in symbiosis and relate these findings to functional changes upon algal / Radiolaria association. We identified compounds from the primary metabolism, vitamins, and specialized metabolites, such as osmolytes, in both investigated systems and putatively assign them to the respective producer. These results pave the way for further investigation of the Radiolaria photosymbiosis, such as attempts to culture them by supplying identified key metabolites, and to support molecular studies by pathway analysis.

## Data availability

The authors declare that all data supporting the findings of this study are available within the paper and its Supplementary files, or public repository. The metabolomics and MSI data have been deposited at the MetaboLights database (Study MTBLS11435).

## Acknowledgements

We thank the Gordon and Betty Moore Foundation for funding within the framework of the Symbiochip project. This work benefited from access to the IMEV, an EMBRC-France and EMBRC-ERIC Site. The authors thank Cyril Debost (IMEV, EMBRC) for the help in the collection of the samples on site. Fluorescent images of Radiolaria were acquired in the microscopy platform PIM (member of MICA microscopy platform). PIM is supported by EMBRC-France, whose French state funds are managed by the ANR within the Investments of the Future program under reference ANR-10-INBS-0. The authors would like to thank Sébastien Schaub (Microscopy Imaging Platform, IMEV, EMBRC) for the technical support in obtaining the light-sheet and confocal images of Radiolaria. CH acknowledges funding by European Union within the framework of European Regional Development Fund (ERDF, Mass Spectrometry Imaging ID 2022FGI0010) CH thanks the Deutsche Forschungsgemeinschaft (DFG, German Research Foundation) for a Leibniz Award. GP and CH thank the Deutsche Forschungsgemeinschaft– Project-ID 239748522 – SFB 1127 ChemBioSys, and the and Germany’s Excellence Strategy—EXC 2051—Project-ID 390713860 (Balance of the Microverse).

## Author contributions

V.N., J.S.M, C.L collected Radiolaria samples; V.N. cultivated the algae, prepared the samples and standards and performed LC-MS experiments and data analysis; V.N. and B.B. prepared samples for MSI experiments; B.B. performed MSI experiments and data analysis; G.P, F.N., C.H., and J.D. acquired funding; V.N. and G.P wrote the main drafts of the manuscript; all authors discussed provided feedback and revisions to the manuscript.

