## Supporting material for "The give and take in photosymbiosis unraveled by metabolomics of Radiolaria"

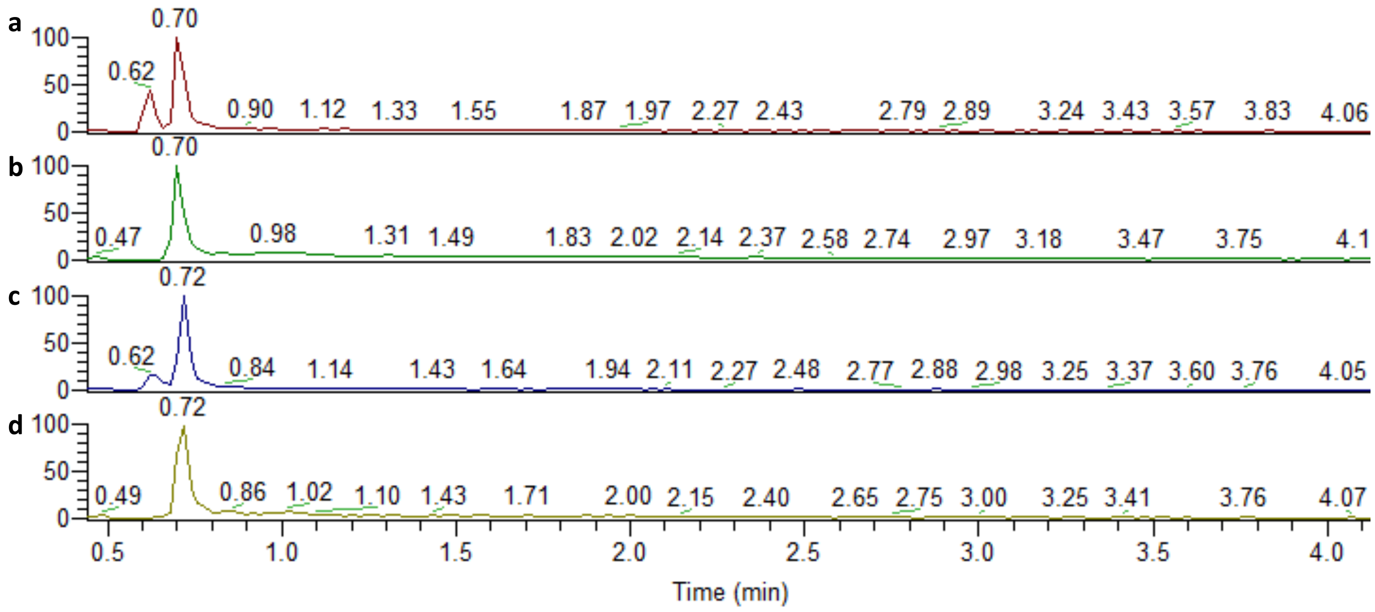


Appendix S1: Extracted ion chromatograms for nicotinamide – *m/z* 123.0553 ± 5ppm (**a, c**), and niacin – *m/z* 124.0394 ± 5ppm (**b, d**) for *Brandtodinium nutricula* cultivated with niacin (**a, b**) and without niacin (**c, d**).

Appendix S2: MALDI-2-MSI of a Collodaria colony section.

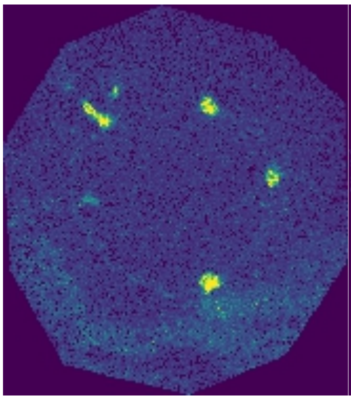

(+) 158.0437 ± 0.0050amu+2.0ppm - Adenine M+Na

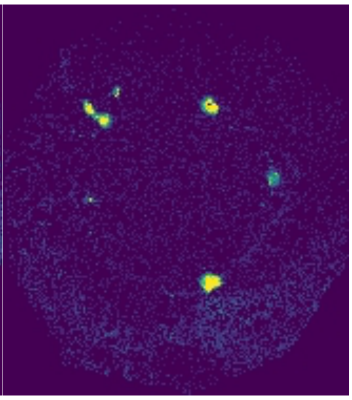

(+) 135.0301 ± 0.0050amu+2.0ppm - Guanine M-$NH_3$+H

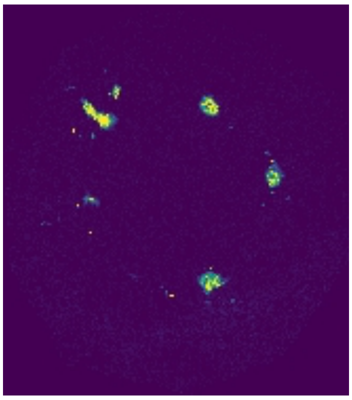

(+) 174.0386 ± 0.0050amu+2.0ppm - Guanine M+Na

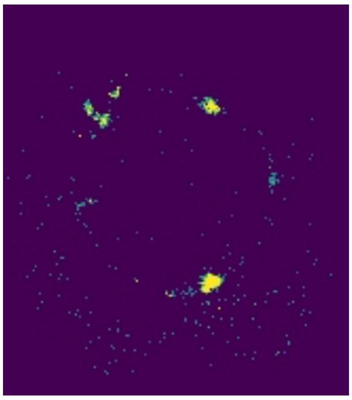

(+) 158.0924 ± 0.0050amu+2.0ppm - Arginine M-$NH_3$+H

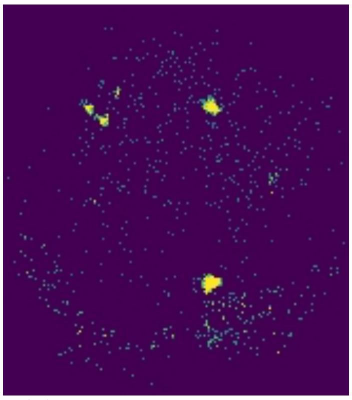

(+) 175.1190 ± 0.0050amu+2.0ppm - Arginine M+H

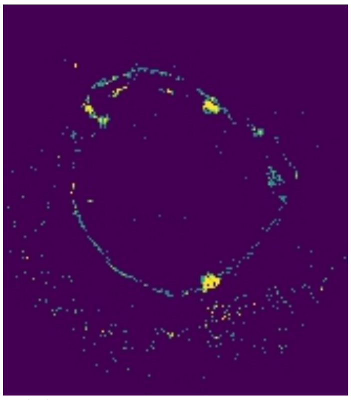

(+) 138.0525 ± 0.0050amu+2.0ppm - Proline M+Na

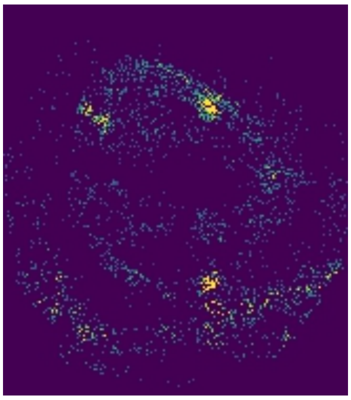

(+) 166.0839 ± 0.0050amu+2.0ppm - Stachydrine M+Na

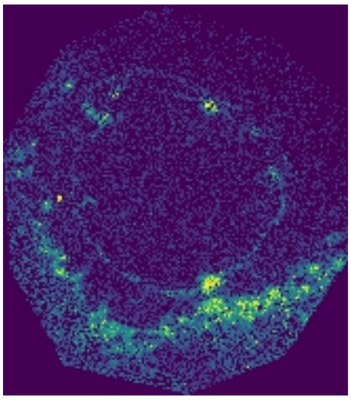

(+) 205.0972 ± 0.0050amu+2.0ppm - Tryptophan M+H

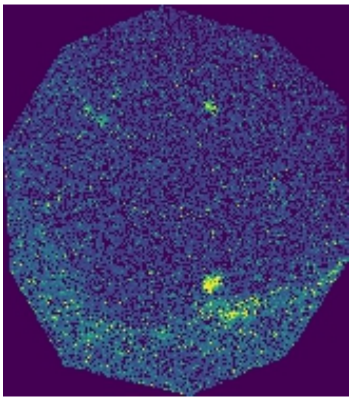

(+) 227.0791 ± 0.0050amu+2.0ppm - Tryptophan M+Na

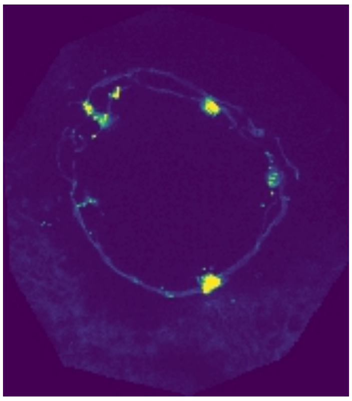

(+) 130.0649 ± 0.0050amu+2.0ppm – $C_9H_8N^{\bullet}$

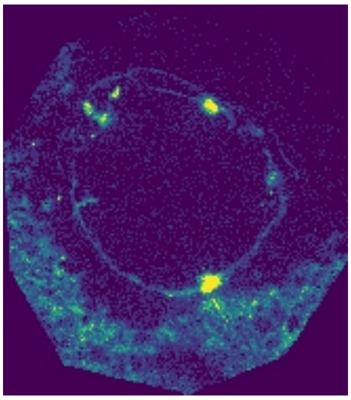

(+) 147.0764 ± 0.0050amu+2.0ppm - Glutamine M+H

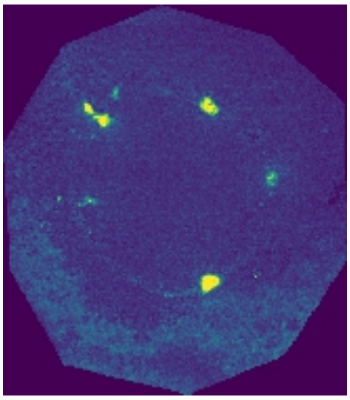

(+) 184.0734 ± 0.0050amu+2.0ppm - γ-butyrobetaine M+K

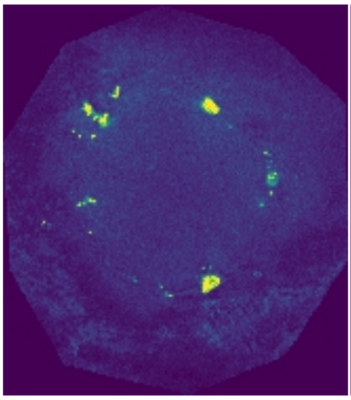

(+) 104.1075 ± 0.0050amu+2.0ppm - Choline $M^+$

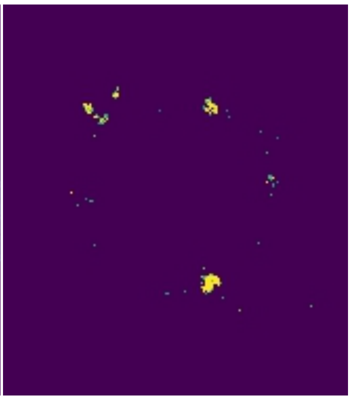

(+) 126.0895 ± 0.0050amu+2.0ppm - Choline M+Na

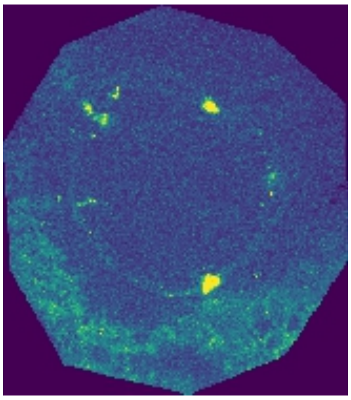

(+) 142.0634 ± 0.0050amu+2.0ppm - Choline M+K

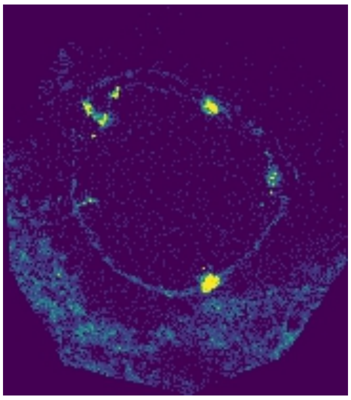

(+) 132.0768 ± 0.0050amu+2.0ppm - Creatine M+H

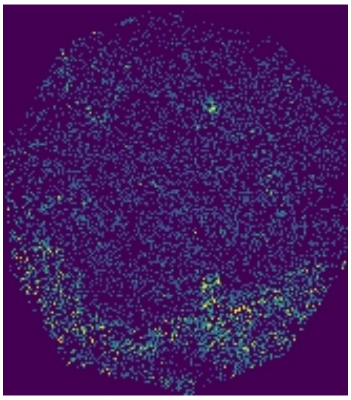

(+) 157.0294 ± 0.0050amu+2.0ppm - DMSP M+Na

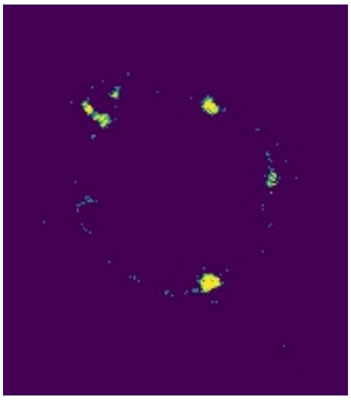

(+) 99.0917 ± 0.0050amu+2.0ppm - Ectoine M-$CO_2$+H

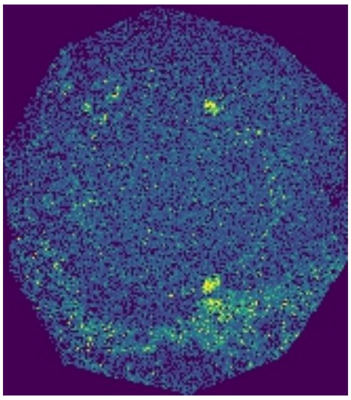

(+) 181.0374 ± 0.0050amu+2.0ppm - Ectoine M+K

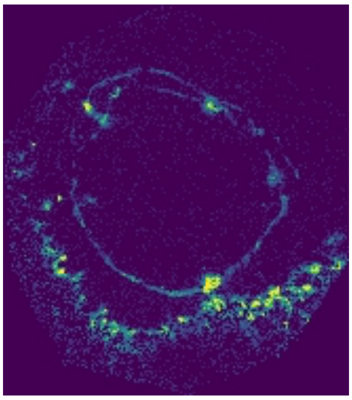

(+) 135.0838 ± 0.0050amu+2.0ppm - Gonyol M-$CO_2$+H

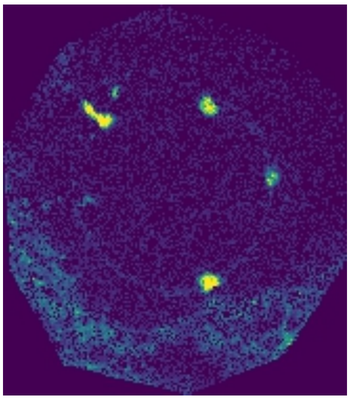

(+) 161.0631 ± 0.0050amu+2.0ppm - Gonyol M-$H_2O$+H

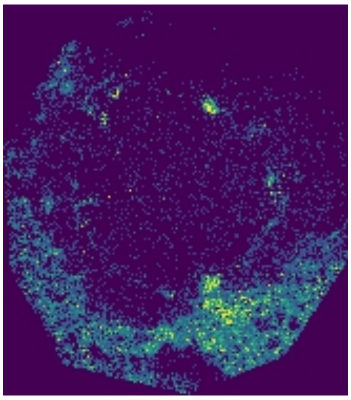

(+) 176.0108 ± 0.0050amu+2.0ppm - Homarine / Trigonelline M+K

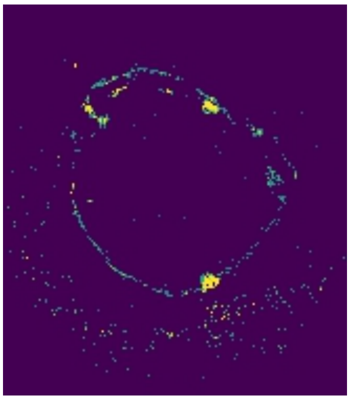

(+) 138.0550 ± 0.0050amu+2.0ppm - Homarine / Trigonelline M+H

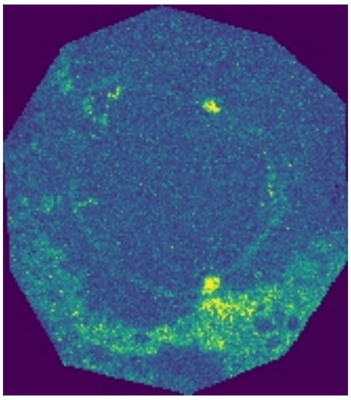

(+) 160.0369 ± 0.0050amu+2.0ppm - Homarine / Trigonelline M+Na

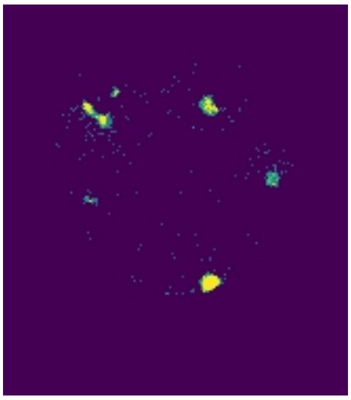

(+) 478.3292 ± 0.0050amu+2.0ppm - 1-palmitoyl-sn-glycero-3-phosphocholine M-$H_2O$+H

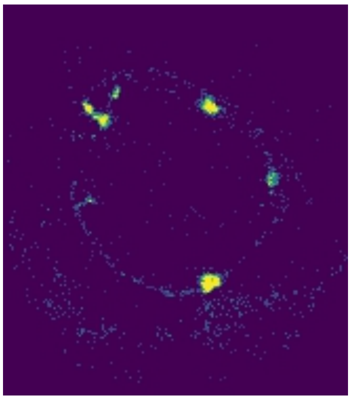

(+) 95.0603 ± 0.0050amu+2.0ppm - urocanic acid M-$CO_2$+H

Appendix S3: Microscopic images

Brightfield microscopy – Image of Collodaria colonies were taken with a stereoscope Leica S AP0 and an INFINITY3-3 camera. Images of Acantharia and the central capsule of Collodaria were taken with a Leica DM IL LED microscope equipped with a Leica DFC280 camera.

Fluorescent microscopy – Fluorescent images of Collodaria colony were taken with SP8 Leica confocal microscope (excitation 638 nm, emission 645–671 nm). Fluorescent image of Acantharia was taken with Viventis LS2 light-sheet microscope (excitation 405 nm, emission 647–800 nm).

Appendix S4: List of standards used for LC-MS measurements.

Ectoine, nicotinate, nicotinamide, GABA betaine hydrochloride, diethanolamine, thiamine, myristoylcarnitine, spermidine, sphingenine from MSMLS, urocanic acid from MSMLS (Sigma Aldrich); betaine, acetylcarnitine hydrochloride, propionylcarnitine, n-dimethylarginine, choline chloride (Sigma); stachydrine (Fluorochem); L-arginine monohydrochloride, L-glutamine, L-leucine, L-phenylalanine, L-proline, L-tyrosine, L-valine, guanine, trigonelline hydrochloride, fucoxanthine, L-isoleucine, L-tryptophane (Fluka analytical); adenosine, pipecolinic acid, hypoxanthine, 1-(2-Hydroxyethyl)piperazine (AlfaAesar); linoleamide (Santa Cruz Biotechnology, Inc.); sulfobetaine (ABCR GmbH); (14:0) 1-myristoyl-sn-glycero-3-phosphocholine, (16:0) 1-palmitoyl-sn-glycero-3-phosphocholine (Avanti Polar Lipids Part of Corda International Plc); adenine (Carl Roth); taurine (Merck KGaA); creatine monohydrate, 3-methyladenine (Thermo Fisher Scientific); phytosphingosine (Tokyo Chemical Industry Co., Ltd); hydroxyproline betaine, β-alanine betaine, alanine betaine (synthetized as described below (Appendix S5); DMSP, homarine, gonyol [1].

Appendix S5: Synthesis of betaines

Since analytical standards for alanine betaine, β-alanine betaine, and hydroxyproline betaine were not available, the betaines were synthesized using a method described in Chen and Benoiton [2] with some modifications. Briefly, 1 g of $KHCO_3$ was dissolved in 20 mL of methanol by stirring for 1 h at 30 °C. Of this solution, 300 µL were added to 3 µg of the corresponding amino acid (alanine, β-alanine, or hydroxyproline). Subsequently, 60 µL of iodomethane were added to each sample, and samples were vortexed for 24 h at room temperature. After that, 100 µL of formic acid were added, samples were dried under vacuum and stored at –20 °C. For LC-HR-MS analysis, samples were dissolved and subsequently diluted 1 : 1000 (*v*:*v*) in either a mixture of methanol: acetonitrile: water (5:9:1, *v*:*v*:*v*) for β-alanine betaine, and hydroxyproline betaine, or in $H_2O$ for alanine betaine. The verification of compounds was done based on their accurate mass.

Appendix S6: LC-MS instrument settings.

MS1 measurements were conducted with the following settings: automatic gain control target – 3E6, the maximum ion injection time – 200 ms, and scan range from 80 to 1200 m/z. Single samples were measured simultaneously with positive and negative modes with resolution 70,000. The order of measurement for samples was randomized.

MS2 measurements were performed for 200 most intense signals among each group of compounds (algal unique compounds, holobiont unique compounds, common compounds) detected with the positive ionization mode. The measurements were performed with positive ionization mode using parallel reaction monitoring with inclusion lists created for each group of compounds. The duration time and runtime were the same as for the MS1 measurement, the settings were as followed: automatic gain control target 2E5, maximum ion injection time 100 ms, three-stepped normalized collision energy 15, 30, 45, scan range from 80 to 1200 m/z, resolution 70,000, in profile mode.

Appendix S7: Sirius settings

Raw files were converted into .mzML format using Proteowizard Suite (proteowizard.sourceforge.net) with vendor peak picking enabled. Putative identification of unknowns was performed depending on the tree fragmentation score [3], and the percentage of CSI:FingerID tool [4]. In case of identification to compound class, the CSI:Finger ID was ≥65%, and Posterior probability of the compound class was ≥90%. The identity of unknowns was confirmed by comparison of retention time and MS2 spectra with chemical standards (accepted difference ≤0.2 min, < 5 ppm).

Appendix S8: MALDI-matrices test

The MALDI-matrices CHCA, DHAP, and DHB (all Bruker, Bremen, Germany) were tested against a panel of 32 compounds, previously identified in the metabolomic experiments. To that purpose, 2 µL of each liquid standard (Appendix S4) were spotted three times on a ground steel MALDI target (Bruker) and dried in a desiccator shuttle. Each cluster of spotted standards was then sprayed with one of the three matrices in a M3+ sprayer (HTXImaging, Chapel Hill NC, USA), using the methods described below (Appendix S9). Mass spectra were acquired in positive ion mode on a timsTOF fleX MALDI-2 mass spectrometer (Bruker) with and without laser post-ionization, after calibrating and tuning the system for optimal detection with red phosphorous (Appendix S10, S11).

Appendix S9: Matrix application methods used for applying MALDI matrices to dried liquid standards on a MALDI target plate and cryosections with the M3+ sprayer.

| Matrix | CHCA | DHB | DHAP |
|---|---|---|---|
| Solvent | 3:1 ACN/H2O + 0.1% FA | 3:1 MeOH/H2O + 0.1% FA | 3:1 ACN/H2O + 0.1% FA |
| Concentration in [mg/mL] | 10 | 50 | 20 |
| Nozzle temp. [°C] | 80 | 90 | 80 |
| Nebulizer gas pressure [psi] | 10 | 10 | 10 |
| FLOW RATE [µl/min] | 100 | 75 | 50 |
| Nozzle velocity [mm/min] | 1250 | 1200 | 1200 |
| Track spacing [mm] | 2 | 2 | 2 |
| Number of layers | 5 | 5 | 5 |
| Nozzle meandering | horizontal and vertical | horizontal and vertical | horizontal and vertical |
| Drying time [s] | 30 | 15 | 30 |

Appendix S10: Measured ion intensities of dried liquid standards acquired with MALDI and MALDI-2 and three different matrices – CHCA, DHAP, and DHB; Each value is based on the integration of 1000 singular spectra, acquired by randomly walking in a 2 mm radius every 10 shots; The liquid standards were spotted and dried prior to the matrix application.

| | MALDI E=50% | | | MALDI2 E=15% | | |
|---|---|---|---|---|---|---|
| Compound\Matrix | DHAP | DHB | CHCA | DHAP | DHB | CHCA |
| Choline | 4954931 | 206788 | 4998204 | 16132439 | 5438562 | 21774266 |
| Proline | 124674 | 421 | 181 | 284214 | 20288 | 48421 |
| Valine | 1011 | 419 | 3483 | 62575 | 8786 | 17031 |
| Betaine | 16780 | 0 | 7262 | 3021204 | 471708 | 3741286 |
| Sulfobetaine | 58138 | 13720 | 1510 | 723903 | 43026 | 165267 |
| Taurine | 2469 | 203 | 15638 | 5388 | 1163 | 116605 |
| Pipecolate | 145809 | 11490 | 66 | 1469786 | 174930 | 80955 |
| Creatine | 707055 | 67819 | 756299 | 511408 | 94601 | 265075 |
| Isoleucine | 2913 | 270 | 551 | 16298 | 4916 | 42721 |
| Leucine | 2798 | 100 | 114 | 47665 | 4473 | 187295 |
| Alanine betaine | 32568 | 5877 | 42316 | 238621 | 34429 | 180726 |
| β-alanine betaine | 234386 | 65736 | 979 | 434678 | 137734 | 1394389 |
| DMSP | 1002478 | 34344 | 2151095 | 2104806 | 254106 | 2384499 |

| | | | | | | |
|---|---|---|---|---|---|---|
| Adenine | 1691795 | 100017 | 2023584 | 7433603 | 999917 | 6942593 |
| Homarine | 70799 | 19135 | 192708 | 613869 | 97916 | 500666 |
| Trigonelline | 1444975 | 379965 | 1831846 | 6285249 | 866113 | 3713875 |
| Ectoine | 5242879 | 159715 | 2299955 | 15570762 | 1338777 | 9005446 |
| Stachydrine | 3404507 | 28297 | 6044 | 15072645 | 1012819 | 8401989 |
| γ-butyrobetaine | 475383 | 62997 | 14575 | 1133828 | 210193 | 2095191 |
| Glutamine | 2779 | 493 | 6925 | 93600 | 14581 | 164229 |
| Guanine | 31751 | 1072 | 32650 | 432217 | 14734 | 236882 |
| Phenylalanine | 6420 | 530 | 13817 | 446214 | 86272 | 970758 |
| Arginine | 240405 | 226463 | 2447430 | 3343689 | 1256281 | 6930484 |
| Gonyol | 11942 | 7286 | 24933 | 29450 | 15912 | 28853 |
| Tyrosine | 17032 | 743 | 3627 | 342991 | 158949 | 994382 |
| n-dimethylarginine | 3675647 | 191024 | 2436833 | 11912504 | 1592001 | 16045597 |
| Tryptophan | 6324 | 337 | 831 | 2075917 | 606857 | 3874878 |
| Propionylcarnitine | 2072645 | 93536 | 3060409 | 5024218 | 522685 | 5912747 |
| Adenosine | 1057315 | 123643 | 52962 | 731699 | 420388 | 227070 |
| Sphingenine | 659 | 686 | 320 | 905 | 2074 | 917 |
| (14:0) 1-myristoyl-sn-glycero-3-phosphocholine | 545799 | 212106 | 892198 | 6723173 | 992205 | 3266230 |
| (16:0) 1-palmitoyl-sn-glycero-3-phosphocholine | 1693255 | 139925 | 1521573 | 5338927 | 1642067 | 4785284 |
| Sum Intensities | 2897832<br>1 | 215515<br>7 | 2484091<br>8 | 10765844<br>5 | 1853946<br>3 | 10449660<br>7 |
| Average Intensity | 905573 | 67349 | 776279 | 3364326 | 579358 | 3265519 |
| Median intensity | 135242 | 16428 | 20286 | 727801 | 166940 | 982570 |

Appendix S11: Bar graph of measured intensities ionizing a panel of analytical standards with MALDI and MALDI-2 and three different matrices CHCA, DHAP, and DHB; The liquid standards were spotted and dried prior to the matrix application.

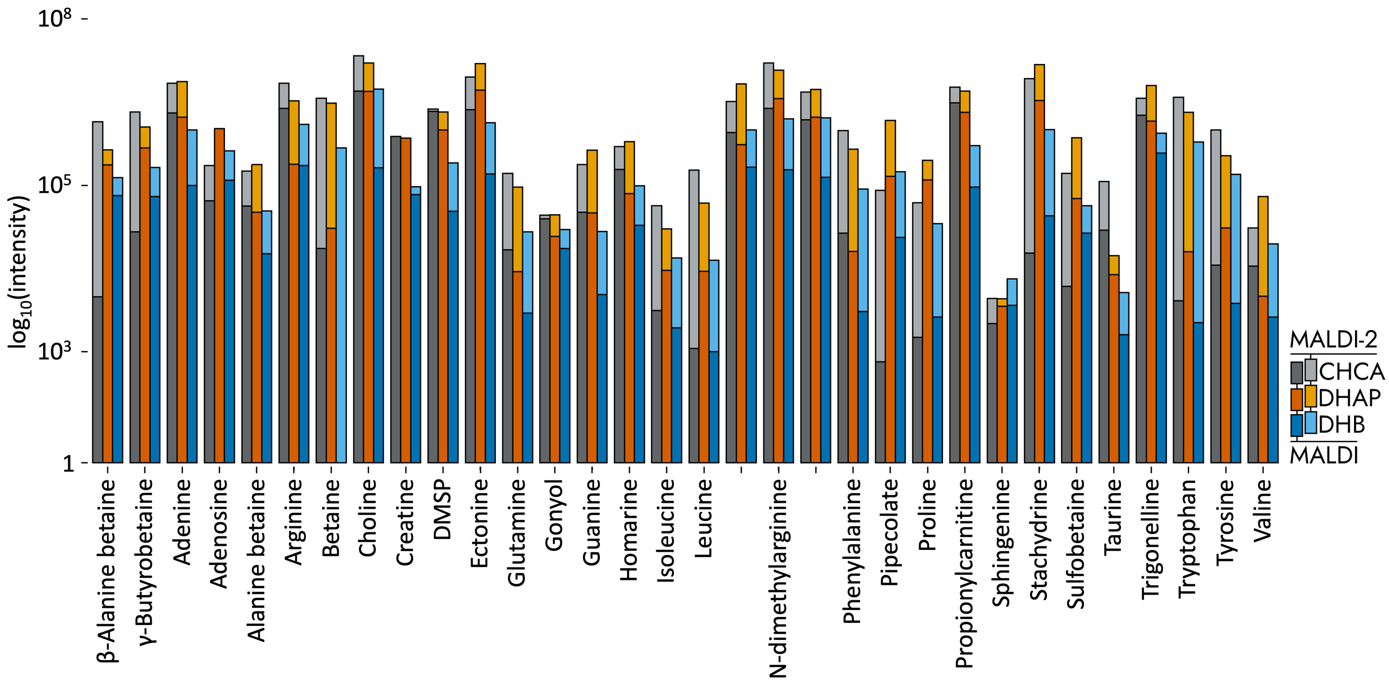


Appendix S12: Possible origin of *N*-(2-hydroxyethyl)piperazine in algal samples.

*N*-(2-hydroxyethyl)piperazine was detected for both cultures of the free-living algae but not in the radiolarian samples. However, since no information about the bioactive role of this molecule could be found in the literature, it was assumed, that the origin of the compound could be the medium that contained 4-(2-hydroxyethyl)-1-piperazineethanesulfonic acid (HEPES) as a buffer. The higher content in the algal samples could be explained by the adsorption of the compounds on the surface of the algae.
